# Hallmarks and metabolic regulation of type 2 activated human lung macrophages

**DOI:** 10.1101/2025.02.26.640352

**Authors:** Amanda J. L. Ridley, Annabel J. Curle, Stefano A. P. Colombo, Joshua J. Hughes, Douglas P. Dyer, Angela Simpson, Maria Feeney, Peter C. Cook, Andrew S. MacDonald

## Abstract

Although human lung macrophages are heterogenous and play key roles during health and disease, the mechanisms that govern their activation and function are unclear, particularly in type 2 settings. Our understanding of how human lung macrophages respond to inflammatory signals have predominantly relied on cell lines or peripheral blood derived cells, which have a limited capacity to reflect the complexity of tissue macrophage responses. Therefore, we isolated macrophages from resected human lung tissue and stimulated them *ex vivo* under type 2 (IL-4, IL-13, or IL-4 + IL-13) or type 1 (IFNγ + LPS) conditions. Human lung macrophages stimulated with IL-4/13, alone or in combination, significantly upregulated expression of the chemokines *CCL17*, *CCL18* and *CCL22,* along with the transglutaminase *TGM2* and the lipoxygenase *ALOX15*. This type 2 activation profile was distinct from LPS + IFNγ activated human lung macrophages, which upregulated *IL6*, *IL8*, *IL1β*, *TNFα* and *CHI3L1* (YKL-40). Further, type 2 activated human lung macrophage products showed differential metabolic reliance for their induction, with IL-4/13 induced *CCL22* being glycolytically controlled, while *ALOX15* was regulated by fatty acid oxidation. These data clarify hallmarks of human lung macrophage activation and polarisation in addition to revealing novel metabolic regulation of type 2 markers.

## Introduction

Lung macrophages are highly specialised, playing a central role in maintaining health and restricting disease (1). Their development, metabolic responsiveness and function are known to be shaped by the unique metabolically challenging lung environment since its available metabolic substrates are tightly controlled (2–4). For instance, the airways are abundant in lipid-rich surfactant while having one of the lowest glucose environments in the body, with levels reported to be less than twelve times that found in the blood (5–7).

Metabolism governs how macrophages respond to signals, including type 1 and type 2 cytokines. With the general principle being that glycolysis tends to be more typical of type 1 macrophages, while lipid metabolism may be more associated with type 2 macrophages (8, 9). Although the emphasis of work on human lung tissue macrophage immunometabolism has so far focused on LPS responsiveness, rather than on type 2 stimuli such as IL-4 and IL-13 (10).

Given limitations on collecting primary human tissue cells (11), our current understanding of human lung macrophage function and immunometabolism during disease predominantly comes from studies using human peripheral blood monocyte derived macrophages (MDMs) *in vitro,* or is extrapolated from murine *in vivo* models (12–14). However, as the tissue environment plays a central role in shaping cellular fate, metabolic responsiveness, and function (Hussell and Bell, 2014; Bain and MacDonald, 2022), it is unlikely that MDMs are representative of lung macrophage function and immunometabolism (8, 9). Similarly, the direct translatability of observations in murine lung macrophages to humans is unclear, with previously defined differences being apparent (15, 16). Such differences likely contribute to why so many drug development programmes fail in translation from preclinical models (17, 18).

There also remains a basic need to define robust markers to identify human tissue macrophages in different activation states, particularly in the context of type 2 (IL-4 and IL-13 driven) inflammation (1, 16, 19). Flow cytometric identification of lung type 2 macrophage activation markers is challenging, since human lung macrophages are highly auto-fluorescent and some commonly used murine markers of type 2 associated macrophages such as *Chil3* have no direct human homologs (20–23). Here, we demonstrate that CCL17, CCL22, ALOX15, CCL18 and TGM2 represent reliable markers of human lung macrophage type 2 activation, and that expression of these markers show distinct metabolic regulation to their type 1 counterparts.

## Results

### Human lung alveolar macrophages display distinct polarization phenotypes following stimulation

To define changes in gene expression and protein secretion in human lung macrophages during polarization, we isolated macrophages by perfusion of human lung tissue obtained from patients undergoing surgical tissue resection with no underlying autoimmune diseases or COPD diagnosis. Macrophages were cultured *ex vivo* under type 2 (IL-4 and/or IL-13) or type 1 (LPS + IFNγ) polarising conditions for 48 h, RNA isolated to quantify gene expression and supernatants collected for measurement of secreted proteins. Macrophages showed distinct profiles of gene expression (Figure 1A). Type 2 polarization resulted in significantly increased expression of *CCL17* (Figure 1B)*, CCL22* (Figure 1C), *TGM2* and *ALOX15,* along with a trend for elevated *CCL18*. In contrast, type 1 stimulation significantly increased expression of *IL6, TNFα*, *IL8, IL1B* and *CHI3L1* (Figure 1A).

**Figure 1.**
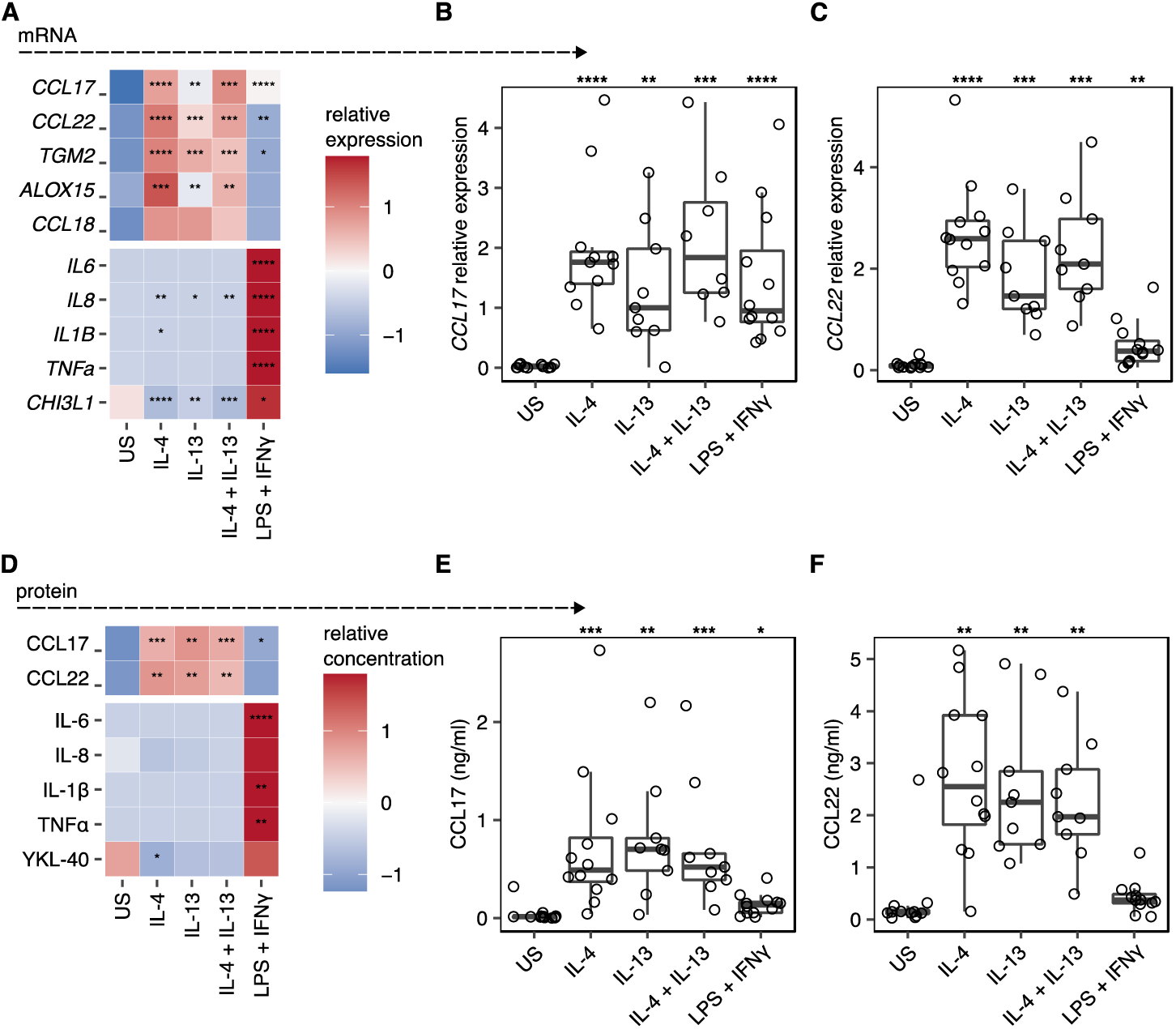
CCL17 is a marker of IL-4 and IL-13 stimulated *ex vivo* lung macrophages. (A-C) Quantification of mRNA expression in human lung macrophages following stimulation with IL-4, IL-13, IL-4 + IL-13, or LPS + IFNγ. Relative gene expression was normalized to *HPRT* using the 2^-ΔCT^ method. US indicates unstimulated controls. (A) Heatmap representation of relative gene expression scaled by row. (B&C) Scatter plots showing the individual sample gene expression values for (B) *CCL17* and (C) *CCL22*. Points represent individual patients. Box plots indicate the median and IQ range, whiskers indicate 1.5x the quartile limit. (D-F) Quantification of protein secretion into the supernatant of macrophages stimulated with IL-4, IL-13, IL-4 + IL-13, or LPS + IFNγ, measured by ELISA. (D) Heatmap showing the relative quantification of different proteins scaled to the row value. (E&F) Scatter plots showing the protein concentration of (E) CCL17 and (F) CCL22. Boxes indicate the median and IQ range, whiskers indicate 1.5x the quartile limit. Statistical values were calculated using Wilcoxon two-sample tests and adjusted for multiple testing using the Holm method. * indicates adjust *P* < 0.05, ** indicates adjust *P* < 0.01, *** indicates adjust *P* < 0.001, **** indicates adjust *P* < 0.0001.

Consistent with our mRNA expression data, we observed significant increases in CCL17 and CCL22 in supernatants of type 2 polarized macrophage cultures, but not type 1 polarised macrophages, which instead demonstrated elevated levels of IL-6, IL-8, IL-1β, TNFα, and YKL-40 (the product of *CHI3L1*) (Figure 1D-F). For the first time, we show that CCL17 represents a robust marker of IL-4 and IL-13 stimulated human lung macrophages, in addition to revealing CCL18 and TGM2 as being associated with IL-4 or IL-13 activated human lung macrophages.

### Type 2 polarization of human lung macrophages is metabolically dependent

The lungs are a metabolically challenging environment, with tightly controlled metabolic substrate availability (2–4). To determine whether human lung macrophages display specific metabolic dependence during type 2/type 1 polarization, we added either 2-DG (a competitive inhibitor of glucose metabolism which blocks the glycolytic pathway (24)), or Eto (an inhibitor of fatty acid oxidation (FAO) (25)) (Figure 2A and Supplemental Figure 1).

**Figure 2.**
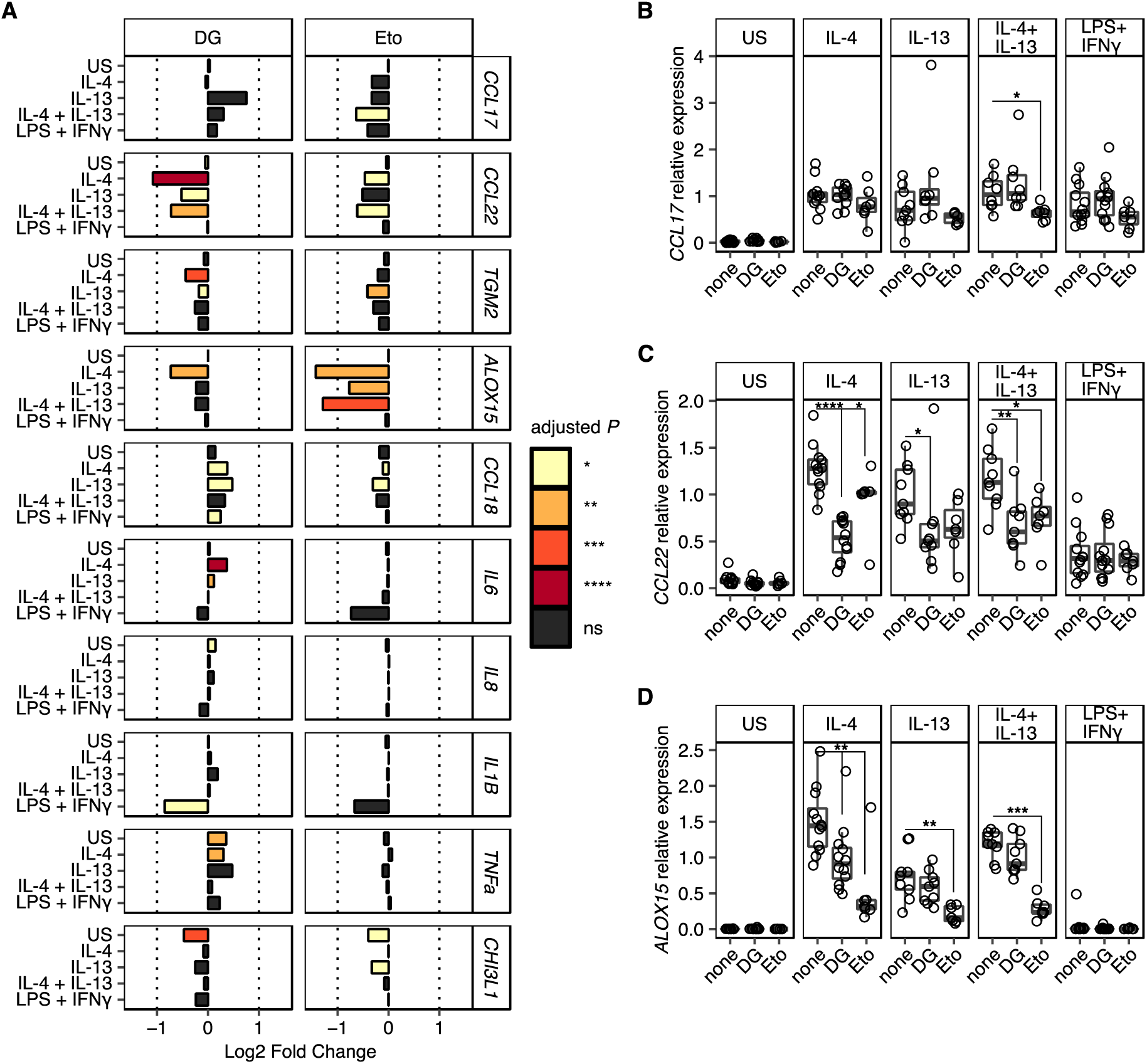
Metabolic dependence in gene expression during type 2 macrophage polarization. Macrophages isolated from resected human lung tissue were stimulated with IL-4, IL-13, IL-4 + IL-13, or LPS + IFNγ and cultured in the presence of metabolic inhibitors, 2-Deoxy-D-glucose (2-DG) or Etomoxir (Eto). Gene expression was quantified by qPCR normalized against HPRT using the 2^-ΔCT^ method. US indicates unstimulated controls. (A) Bar chart representing Log2 Fold Change in gene expression for each gene and stimulation condition. Fold change is expressed relative to the matched culture condition without metabolic inhibitor. Bar colour indicates the level of statistical significance. (B-D) Scatter plots showing relative expression of (B) *CCL17,* (C) *CCL22,* and (D) *ALOX15* under different stimulation conditions with (2-DG & Eto) or without (none) metabolic inhibitors. Boxes indicate the median and IQ range, whiskers indicate 1.5x the quartile limit. Statistical values were calculated using Wilcoxon two-sample tests and adjusted for multiple testing using the Holm method. * indicates adjust *P* < 0.05, ** indicates adjust *P* < 0.01, *** indicates adjust *P* < 0.001, **** indicates adjust *P* < 0.0001.

Culture of human lung macrophages with 2-DG resulted in a significant reduction in the expression of *CCL22* during IL-4-induced type 2 polarization (Figure. 2A and 2C). Secretion of CCL22 into culture supernatants was also impaired under these conditions, although this only reached statistical significance in the IL-4 stimulated condition (Supplemental Figure 1). Additionally, a small but significant reduction in *ALOX15* expression was observed in the IL-4 stimulation condition only (Figure 2A & 2D). In contrast, Eto caused a significant reduction in *ALOX15* expression under all type 2 polarization conditions, with smaller reductions in *CCL17, CCL22, TGM2 and CCL18* expression, which only reach statistical significance under specific type 2 stimulation conditions (Figure 2A and 2B). Consistent with previous observations (26, 27) we observed a significant decrease in *IL-1β* expression during type 1 polarization following addition of 2-DG (Figure 2A). Similarly, type 1 polarised human lung macrophages cultured with Eto showed a strong trend for reduced expression of *IL-1β* and *IL-6* (Figure 2A).

Together, these data promote key roles for both glycolysis and FAO in the polarization of macrophages in a type 2 context, in a marker-specific manner. Specifically, whilst there was a lack of reliance on glycolysis for expression of *CCL17* by type 2 activated human lung macrophages, there was a shared reliance on glycolysis and FAO for *CCL22* expression. Together, these data support a model in which expression of different genes stimulated by type 2 cytokines are dependent on distinct metabolic pathways.

## Discussion

Despite murine models being widely used to study activation and polarization of macrophages, clear markers for identification of type 2 (IL-4 and IL-13) activated human macrophages are lacking. This is mainly since translation of murine data to humans can be problematic, as many macrophage type 2 markers identified in mice (e.g., Chil3/Ym1) ((20) have no direct human homologue, or such markers are differentially regulated in humans (e.g., Arginase 1) (23, 28), or the gene exists in humans but not mice (e.g., *CCL18*) (23, 28). These fundamental differences between mouse and human macrophages highlight the requirement for more markers common to both species to be identified, to enable more meaningful translational studies into macrophages during type 2 inflammatory diseases. At the same time, much of what we currently know about macrophage activation and function is reliant on use of model murine or human macrophages *in vitro*, such as BMDMs or MDMs, rather than those isolated *ex vivo* from relevant tissues.

Our work builds upon and extends previous limited human lung tissue macrophage studies to comprehensively identify a suite of markers - CCL17, CCL22, TGM2, ALOX15 and CCL18 - as hallmarks of type 2 (IL-4 and IL-13) activation of human pulmonary macrophages *ex vivo*. Here, we demonstrate for the first time that CCL17 represents a robust marker of IL-4 and IL-13 stimulated human lung macrophages and confirm previous suggestions that CCL22 may also represent a type 2 human lung macrophage marker (29). Together, our data are in line with previous studies using IL-4 and IL-13 polarised human MDMs (30, 31) and murine BMDMs (32), that suggested that CCL17 (Figure 1B and E) and CCL22 (Figure 1C and F) represent type 2 macrophage markers common to both species.

Other type 2 activation markers we validated for human lung macrophages included ALOX15, CCL18 and TGM2, confirming ALOX15 as a consistent marker of type 2 activated human lung macrophages (29), while revealing CCL18 and TGM2 as being associated with IL-4 or IL-13 activated human lung macrophages (Figure 1A). This supports previous suggestions that isolated human BAL fluid macrophages (33) and human MDMs can express CCL18 in response to IL-4 or IL-13 (31), with the latter also upregulating TGM2 (30, 34, 35).

In addition to increasing clarity surrounding type 2 human tissue macrophage markers, we have also addressed how these cells are governed metabolically. Of note, human lung macrophage expression of the type 2 marker CCL22 following IL-4 and IL-13 stimulation was strikingly reduced upon addition of the glycolytic inhibitor 2-DG (Figure 2C), which strongly contrasted the limited impact of 2-DG on CCL17 expression (Figure 2B). This reflects the fact that, despite CCL17 and CCL22 being known to share the same G-protein coupled receptor CCR4 (36, 37), they are distinct in terms of their structure (36, 38), receptor binding affinity (39), downstream signalling (40) and expression profiles (41–43). Further, this direct association between glycolytic metabolism and CCL22 production supports previous studies which have reported that diet can influence circulating levels of CCL22 and that its production in response to IL-4 by murine BMDMs is reliant on glutamine (44, 45). Our data also demonstrate for the first time that IL-4 induced *ALOX15* mRNA expression by human lung macrophages is controlled by glycolysis and FAO (Figure 2D). Broadly, these data extend previous observations using murine BMDMs and imply that human lung macrophage responsiveness to IL-4 and IL-13 is regulated by both glycolysis and FAO (46–48).

It has been previously reported that FAO is dispensable for human MDM IL-4 responsiveness but not murine MDMs (12, 49). Consistent with this, we demonstrate that human lung macrophage expression of CCL17, CCL22, TGM2 and CCL18 in response to IL-4 and IL-13 was not significantly affected by addition of the FAO inhibitor Eto (Figure 2A). Notably, we did observe a trend for reduced CCL17 and CCL22 expression in this context, indicating a potential role for FAO (Figure 2B and C). Our results also demonstrate that expression of the type 2 marker ALOX15 by human lung macrophages cultured with IL-4 and IL-13 was significantly reduced following addition of Eto (Figure 2C), indicating a role for FAO. Looking at the type 2 markers on an individual basis, it is striking that they appear to be distinctly metabolically programmed, with this level of fine tuning likely to be relevant in the *in vivo* context.

Similarly to human MDMs (Bonneh-Barkay *et al.*, 2012), we have found that YKL-40, the human equivalent of the type 2 murine macrophage marker Ym1 (Raes *et al.*, 2005), was upregulated by human lung macrophages in response to LPS, rather than IL-4 or IL-13 (Figure 1D). However, a role for YKL-40 during type 2 inflammatory disease cannot be ruled out as it has been reported to be elevated in the serum of asthmatics and associated with increased asthma severity (50, 51).

Our data confirm previous studies using human MDMs (30), human airway macrophages (10) and lung macrophages (52, 53) that IL-6, IL-8, IL-1β and TNFα represent type 1 activation markers (Figure 1A and D). We found that, despite displaying a robust LPS and IFNγ response (as evidenced by upregulated IL-6, IL-8, IL-1β, TNFα and YKL-40), addition of the glycolytic inhibitor 2-DG had no significant effect on human lung macrophage expression (Figure 2A) or secretion (Supplemental Figure 1) of these markers. Although not significant, our data suggests that glycolysis may play a role in regulating IL-1β expression and YKL-40 secretion in type 1 stimulated human lung macrophages, since there was a strong trend for reduced expression of these markers following addition of 2-DG (Figure 2A and Supplemental Figure 1). Whilst the lack of significant impact of 2-DG on these type 1 markers somewhat contradicts murine studies showing that glycolysis controls murine BMDMs responding to LPS (26, 54), it supports work indicating that glycolysis may be dispensable for LPS activated human MDMs (14, 44). Consistent with this, one of the few immunometabolism studies to use human BAL fluid macrophages found that they failed to display metabolic reprogramming towards glycolysis after LPS stimulation (10), contrary to murine BMDMs (49, 55) and human MDMs (46). Notably, like 2-DG, there was also a strong trend for reduced expression of *IL1β* along with *IL6* mRNA following addition of Eto (Figure 2A), although the reverse was seen in terms of secretion (Supplemental Figure 1).

We and others have previously highlighted the use of tissue macrophages for metabolic profiling, noting that BMDMs and MDMs *in vitro* behave differently and exhibit different metabolic phenotypes to murine or human tissue macrophages *ex vivo* (54, 56). Building on these earlier observations, our current data further demonstrate the importance of using human tissue macrophages to address paradigms in metabolic requirements for activation and function that have previously been constructed using MDMs or murine macrophages. We show that human tissue macrophage metabolic programming is not as simple as once thought, and requires further delineation, as glycolysis and FAO likely play complementary roles in specific aspects of both type 1 and type 2 human macrophage polarisation. Together, our data highlights the critical role of the tissue environment in controlling human macrophage responsiveness to cytokines, a principle that should be considered when interpreting data generated in murine studies or using model human macrophages, given that impaired macrophage metabolism is associated with many human diseases (57–59).

Lung macrophage heterogeneity is complex, consisting of both alveolar and interstitial subsets (60–62), and we and others have previously demonstrated that murine alveolar macrophages show differential metabolism and responsiveness to polarising stimuli than interstitial macrophages (56, 63). Thus, we anticipate that IL-4 and IL-13 induced CCL17 and CCL22 expression by human alveolar and interstitial macrophages may differ, along with their metabolic profiles. This underscores the need for studies to unequivocally distinguish human airway alveolar macrophages from tissue interstitial macrophages, to be able to address such possibilities experimentally.

In summary, our data show that CCL17 and CCL22 along with TGM2, ALOX15 and CCL18, represent reliable markers that can be used to identify and distinguish type 2 (IL-4 and IL-13) from type 1 (LPS and IFNγ) activated human lung macrophages. Further, these novel data increase our understanding of how metabolism governs pulmonary macrophage responsiveness and activation, since we reveal that type 2 macrophage markers (including CCL22 and ALOX15) are significantly regulated by glycolysis and FAO, respectively. Our data emphasise the importance for studies on human tissue macrophages *ex vivo*, to reduce reliance on *in vitro* assays and murine models that may fail to represent the human lung environment. We suggest that CCL17 and CCL22 represent useful human type 2 tissue macrophage markers that may provide potential therapeutic targets for future treatments targeting such mediators or the metabolic pathways that control them in pulmonary type 2 associated inflammatory lung disease.

## Methods

### Sex as a biological variable

In this study, sex was not considered as a biological variable.

### Human subjects and samples

All adult human donors gave written informed consent in accordance with the principles expressed in the Declaration of Helsinki and were recruited into the Manchester, Allergy, Respiratory and Thoracic Surgery (ManARTS) Biobank (located in the Northwest Lung Research Centre at the University Hospital of South Manchester NHS Foundation, Wythenshawe). The research protocol was approved by the National Research Ethics Service (NRES) Committee Northwest – Haydock, ethics approval 15/NW/0409). The ManARTS Biobank collected surgical lung samples from lung cancer patients that had undergone surgical lung tumor resection. The regions of lung marginal tissue samples collected for the experiments described in this study were taken by a histopathologist at a set distance from the lung tumor (>6 cm) during resection and thus deemed non-cancerous for research purposes. Lung resection sample exclusion criteria for this study included patients with Rheumatoid Arthritis, a known history of peripheral arterial disease, sarcoidosis, ankylosing spondylitis, Crohn’s disease, autoimmune diseases (e.g. inflammatory bowel disease, multiple sclerosis and type 1 diabetes), conditions requiring immunomodulatory therapy (e.g. Methotrexate and Azathioprine), a history of tuberculosis and chronic obstructive pulmonary disease (COPD) or emphysema diagnosis. A total of twelve donors aged between 49 and 82 years old were used for the experiments detailed in this study. Their patient demographic and clinical information is listed in **Table 1** along with the patient’s medication history in Table 2.

**Table 1.** Demographic and clinical data for patients who donated lung samples denoted in this study.

| No. | Ex- vs never-smoker | Co-morbidities | Age (years) | Gender (M/F) | FEV1 | FVC | Ratio (%) |
| --- | --- | --- | --- | --- | --- | --- | --- |
| 1 | Never | None | 63 | F | 2.42 (112.7%) | 2.95 (115.4%) | 81.8 |
| 2 | Never | None | 56 | M | 2.36 (71%) | 3.55 (85%) | 66.4 |
| 3 | Never | Previous breast cancer<br>Type II diabetes | 73 | F | 1.81 (104%) | 2.33 (110%) | 78 |
| 4 | Ex | Hypercholesterolaemia<br>Hypertension<br>Atrial fibrillation | 78 | F | 1.94 (105%) | 2.69 (119%) | 72.5 |
| 5 | Ex | Hypercholesterolaemia<br>Hypertension<br>Osteoarthritis<br>Angina<br>Abdominal aortic aneurism<br>B cell carcinoma | 82 | M | 3.12 (121%) | 3.83 (109%) | 81.4 |
| 6 | Ex | Hypertension<br>Deep vein thrombosis | 63 | M | 2.17 (67%) | 2.98 (72%) | 72.8 |
| 7 | Ex | Benign prostatic hyperplasia<br>Hypertension<br>Osteoarthritis<br>Transient ischemic attack | 82 | M | 1.84 (70%) | 2.93 (82%) | 63 |
| 8 | Ex | Type II diabetes<br>Previous colon cancer<br>Deep vein thrombosis | 82 | M | 2.87 (104%) | 4.13 (110%) | 69 |
| 9 | Ex | Previous myocardial infarction | 69 | M | 2.66 (85%) | 4.63 (114%) | 57.5 |
| 10 | Ex | Hypercholesterolaemia<br>Pulmonary fibrosis<br>Hypertension | 74 | M | 2.84 (97%) | 3.83 (99%) | 74 |
| 11 | Ex | Mild obstructive airway disease<br>Mild asthma | 70 | F | 1.60 (73.9%) | 2.3 (88.4%) | 69.5 |
| 12 | Ex | Gastroesophageal reflux disease<br>Mild asthma | 49 | F | 2.58 (105%) | 3.5 (122%) | 74 |

The age stated refers to the patients age at the time of lung resection. M = male; F = female; FEV1 = forced expiratory volume in one second; FVC = forced vital capacity; Ratio = FEV1/FVC. Ratio <70% is considered abnormal and may be indicative of COPD.

**Table 2.**
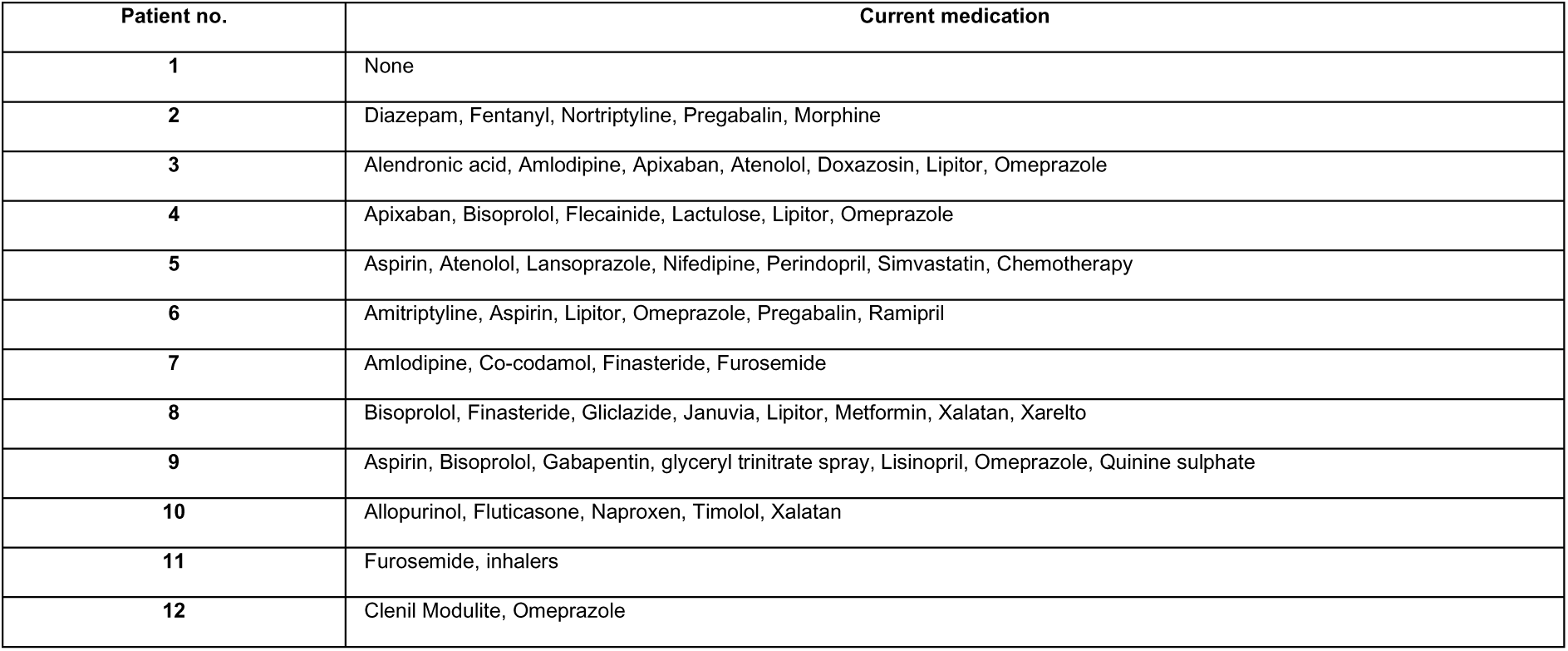
Medication history for patients who donated lung samples denoted in this study.

### Human lung macrophage isolation

Human lung macrophages were isolated from lung tissue resection samples using perfusion and plastic adherence (64). The airways of the resected lung tissue were perfused with sterile 1X PBS using a syringe attached to a 21-gauge needle, 40 mm in length (BD Microlance) until the lung tissue appeared anemic in colour. The perfusate was then centrifuged at 400 x *g* for 10 min at 4 °C (Heraeus Megafuge 40R) and pelleted cells resuspended in sterile 10 ml Roswell Park Memorial Institute (RPMI)-1640 media before being slowly overlaid onto 10 ml Ficoll-Paque Plus (GE Healthcare) using a Pasteur pipette. The cells and Ficoll were then centrifuged at 400 x *g* with reduced acceleration and without brakes for 30 min at RT. After centrifugation, the peripheral blood mononuclear cell (PBMC) layer was collected and washed with 20 ml RPMI-1640 media. The isolated cells were centrifuged at 400 x *g* for 10 min at 4°C before the pelleted cells were resuspended in prewarmed RPMI-1640 media supplemented with 2 mM L-Glutamine (Sigma-Aldrich), 100 U/ml penicillin (Sigma-Aldrich), 100 μg/ml streptomycin (Sigma-Aldrich) and 10% foetal calf serum (FCS) until manually counted based on size and morphology using a haemocytometer, as described previously (65).

### *In vitro* cell culture

#### Human lung macrophage culture with type 2 and type 1 stimuli, and metabolic inhibitors

Cells were resuspended in RPMI-1640 media containing 2 mM L-Glutamine, 100 U/ml penicillin, 100 μg/ml streptomycin and 10% FCS at a density of 4 x 10^5^ cells/ml. Next, 40,000 cells in 100 μl were plated per well in a 96-well tissue culture treated, non-pyrogenic, polystyrene flat-bottomed culture plate with lid (Corning) and allowed to adhere for 1 h at 37°C in a humidified environment of 5% CO2. The total cell counts were based on large cells to ensure enough macrophages were plated and allowed to adhere, with the average purity of macrophages (CD45^+^CD163^+^CD64^+^) after adherence, as determined by flow cytometry, being 75%. After incubation and, adherence, cells were washed three times with 200 μl 1X PBS per well before 100 µl of fresh supplemented RPMI-1640 media was added per well. Next, either 50 μl supplemented RPMI-1640 media alone or containing 4 mM 2-Deoxy-D-glucose (2-DG) (RPMI containing 1 mM final concentration) (Sigma-Aldrich) or 50 μl 800 µM etomoxir (Eto) (200 µM final concentration) (Sigma-Aldrich) was added per well for 30 min at 37 °C, 5% CO2. After incubation, cells were stimulated with either 50 μl supplemented RPMI-1640 media alone or containing 50 μl 80 ng/ml human rIL-4 (20 ng/ml final concentration) (PeproTech) or 80 ng/ml human rIL-13 (20 ng/ml final concentration) (PeproTech) or 80 ng/ml LPS (20 ng/ml final concentration) (Sigma-Aldrich) plus 80 ng/ml IFNγ (20 ng/ml final concentration) (PeproTech) for 48 h at 37 °C, 5% CO2. After 48 h supernatants were harvested and stored at −80 °C for future secreted protein analysis whilst cells were lysed in 150 μl RLT buffer with 1.5 μl 2-ME and stored at −80 °C prior to RNA extraction.

#### ELISA

To quantify protein concentrations in human lung macrophage culture supernatants, ELISAs with paired antibodies and recombinant standards were performed (see **Table 3**). Capture antibodies were coated onto 96 well flat bottom non tissue culture plates (NUNC-Immuno Plate, Thermo Fisher Scientific) in 50 µl of 1X PBS overnight at 4 °C. Plates were then washed four times with 200 µl wash buffer (1X PBS containing 0.05% Tween-20) before being blocked using 50 µl 1% bovine serum albumin (BSA) (Sigma-Aldrich) diluted with 1X PBS for 90 min at 37 °C. After blocking, supernatants in duplicate or triplicate at the required dilution along with doubling dilutions of recombinant protein standards in a volume of 50 µl wash buffer were added. Samples were then incubated overnight at 4 °C. Plates were then washed four times with 200 µl wash buffer before secondary detection antibodies were added in 50 µl wash buffer and allowed to bind for 1 h at 37 °C. Plates were washed a further four times with 200 µl wash buffer before streptavidin-horseradish peroxidase (R&D Systems) diluted 1:200 in wash buffer was added to plates in 50 µl and incubated at 37 °C for 1 h. After incubation, plates were washed eight times with 200 µl wash buffer before 100 µl of colorimetric substrate of peroxidase, 3,3’,5,5’-Tetramethylbenzidine (TMB) (Sigma-Aldrich) was added to each well. Following development, the reaction was stopped by addition of 0.18 M sulfuric acid (Sigma-Aldrich). Plates were read at 450 nm, with 570 nm as the reference wavelength, using a plate reader (Tecan, Infinite M200 PRO).

**Table 3.** ELISA reagents.

| Specificity | Conjugation | Clone | Usage concentration | Manufacturer |
| --- | --- | --- | --- | --- |
| CCL17 (TARC) | Purified | Poly5230 | 0.5 µg/ml | Biolegend |
| CCL22 (MDC) | Purified | - | 2 µg/ml | R&D DuoSet |
| IL-6 | Purified | MQ2-13A5 | 2.5 µg/ml | Biolegend |
| IL-8 | Purified | BH0814 | 2.5 µg/ml | Biolegend |
| TNFα | Purified | Mab1 | 1 µg/ml | Biolegend |
| YKL-40 | Purified | - | 1 µg/ml | R&D DuoSet |
| IL-1β | Purified | - | 4 µg/ml | R&D DuoSet |
| CCL17 (TARC) | Recombinant |  | 1 ng/ml | Biolegend |
| CCL22 (MDC) | Recombinant |  | 1 ng/ml | R&D DuoSet |
| IL-6 | Recombinant |  | 10 ng/ml | Biolegend |
| IL-8 | Recombinant |  | 100 ng/ml | Biolegend |
| TNFα | Recombinant |  | 100 ng/ml | Biolegend |
| YKL-40 | Recombinant |  | 2 ng/ml | R&D DuoSet |
| IL-1β | Recombinant |  | 0.25 ng/ml | R&D DuoSet |
| CCL17 (TARC) | Biotin | Poly5230 | 0.5 µg/ml | Biolegend |
| CCL22 (MDC) | Biotin | - | 0.05 µg/ml | R&D DuoSet |
| IL-6 | Biotin | MQ2-39C3 | 2.5 µg/ml | Biolegend |
| IL-8 | Biotin | BH0840 | 2.5 µg/ml | Biolegend |
| TNFα | Biotin | MAb11 | 0.5 µg/ml | Biolegend |
| YKL-40 | Biotin | - | 0.1 µg/ml | R&D DuoSet |
| IL-1β | Biotin | - | 0.15 µg/ml | R&D DuoSet |

#### RNA extraction and qPCR

Cells post culture were lysed with 150 µl RLT lysis buffer containing 1.5 μl 2-ME before RNA was isolated using RNeasy Micro kits (Qiagen) according to the manufacturer’s instructions. After RNA extraction, cDNA was generated from sorted cells using SuperScript-III (Thermo Fisher Scientific) and Oligo-dT (12-18 primer, Thermo Fisher Scientific). Relative quantification of genes of interest was performed by qPCR analysis with a QuantStudio 12K Flex Real-Time PCR instrument (Thermo Fisher Scientific) using QuantStudio software (Thermo Fisher Scientific) and SYBR Green master mix (Thermo Fisher Scientific). Five serial 1:4 dilutions of a positive control sample of pooled cDNA were used to create standard curves. Gene expression (Ct values) was normalized to Hypoxanthine Phosphoribosyltransferase (*HPRT*). Human primer sequences were checked using the NCBI Basic Local Alignment Search Tool (BLAST) and ordered from Sigma-Aldrich (Germany). Primer pairs for sequences with length between 80-150 were selected (Table 4).

**Table 4.**
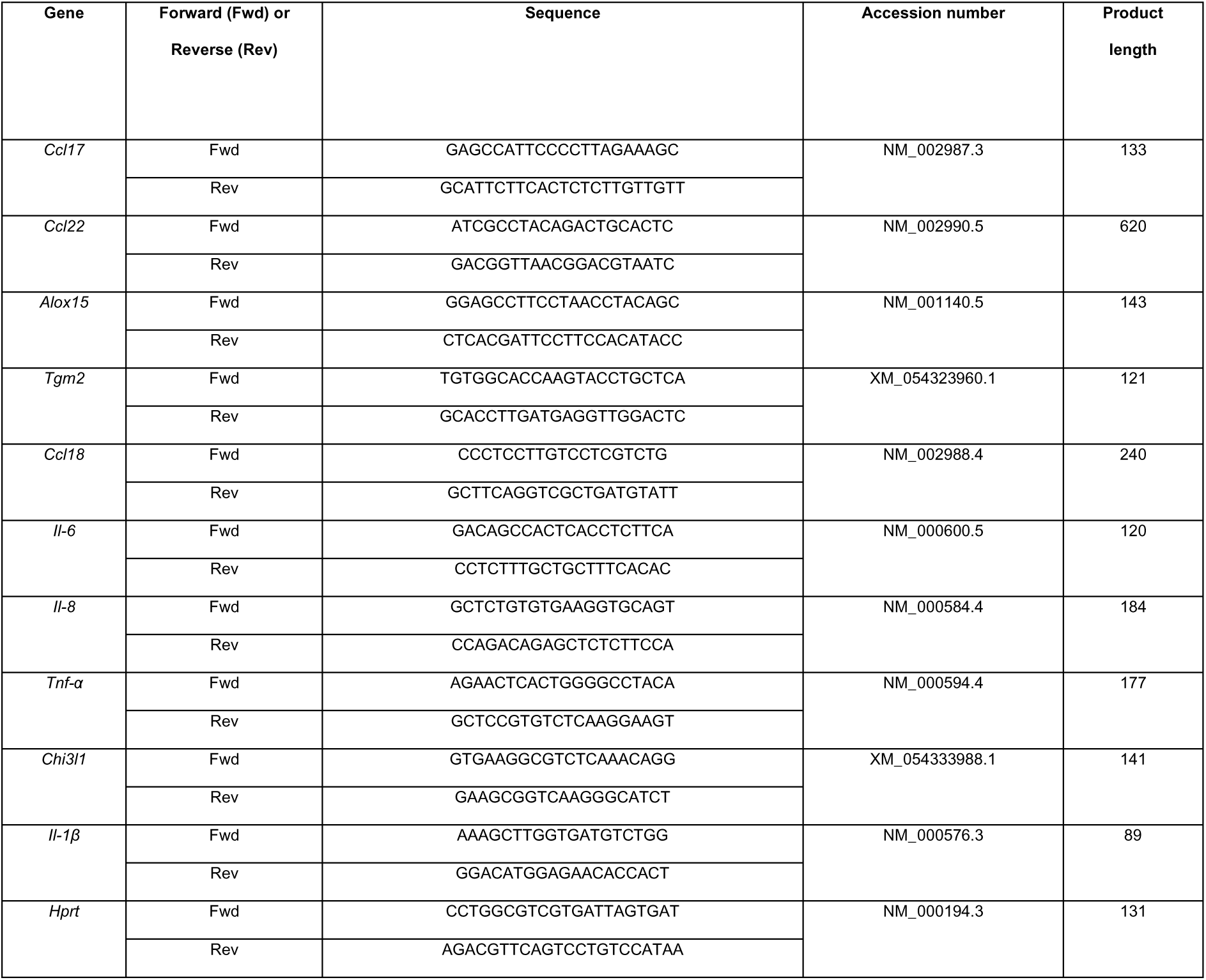
Primers.

#### Statistics

Statistical analysis was performed using R (version 4.3) in RStudio (version 2023.09.1+494). Pairwise comparisons statistical comparisons were calculated by Wilcoxon two-sample tests using the rstatix package (version 0.7.2). *P* values from Wilcoxon comparisons were adjusted for multiple testing using the Holm method. All *P* values are reported as the Holm-adjusted *P* value. An adjusted *P* value less than 0.05 was considered significant. Heatmaps were produced by calculating the geometric mean for each group/condition and plotted using the with tile colours scaled by row. All graphs were produced using the ggplot2 package (version 3.4.1).

#### Study approval

The study was approved under the National Research Ethics Service Committee; North West – Haydock ethics reference 20/NW/0302. Informed consent was obtained from all donors included in this study.

## Data availability

All data needed to evaluate the conclusions in the paper are present in the paper or the Supplemental Materials.

## Author contributions

A.J.L.R., and A.C. performed experiments. S.A.P.C. performed data analyses and prepared figures. A.J.L.R., and A.S.M contributed significantly to design of the study and wrote the manuscript, with constructive input from S.A.P.C., D.P.D., J.J.H., M.F., and P.C.C. A.S. contributed patient resources for the study. All authors contributed to the article and approved the submitted version.

## Acknowledgements

This report is independent research supported by the North West Lung Centre Charity at Manchester University NHS Foundation Trust. The views expressed in this publication are those of the author(s) and not necessarily those of the NHS, the North West Lung Centre Charity or the Department of Health. The authors would like to acknowledge the Manchester Allergy, Respiratory and Thoracic Surgery Biobank and the North West Lung Centre Charity for supporting this project. In addition, we would like to thank the study participants for their contribution. A.J.L.R was supported by a MRC CASE studentship (with GSK) (MR/N013751/1). A.S.M was supported by funding from GSK, the Lydia Becker Institute and the MRC (MR/W018748/1).

