## Supplemental for "Hallmarks and metabolic regulation of type 2 activated human lung macrophages"

**Supplemental material**

**Figures**

**Supplemental Figure 1. Macrophage protein production in the presence of metabolic inhibitors.**

Macrophages isolated from resected human lung tissue were stimulated with IL-4, IL-13, IL-4 + IL-13, or LPS + IFNγ and cultured in the presence of metabolic inhibitors, 2-Deoxy-D-glucose (2-DG) or Etomoxir (Eto). The culture supernatant was collected, and the presence of secreted proteins was quantified by ELISA. Bar chart representing Log2 Fold Change in protein concentration for each protein and stimulation condition. Fold change is expressed relative to the matched culture condition without metabolic inhibitor. Bar colour indicates the level of statistical significance. * indicates adjust *P* < 0.05, ** indicates adjust *P* < 0.01.
